# PoweREST: Statistical Power Estimation for Spatial Transcriptomics Experiments to Detect Differentially Expressed Genes Between Two Conditions

**DOI:** 10.1101/2024.08.30.610564

**Authors:** Lan Shui, Anirban Maitra, Ying Yuan, Ken Lau, Harsimran Kaur, Liang Li, Ziyi Li, the Translational and Basic Science Research in Early Lesions (TBEL) Program

## Abstract

Recent advancements in Spatial Transcriptomics (ST) have significantly enhanced biological research in various domains. However, the high cost of current ST data generation techniques restricts its application in large-scale population studies. Consequently, there is a pressing need to maximize the use of available resources to achieve robust statistical power. One fundamental question in ST analysis is to detect differentially expressed genes (DEGs) among different conditions using ST data. Such DEG analysis is often performed but the associated power calculation is rarely discussed in the literature. To address this gap, we introduce, PoweREST (https://github.com/lanshui98/PoweREST), a power estimation tool designed to support power calculation of DEG detection with 10X Genomics Visium data. PoweREST enables power estimation both before any ST experiments or after preliminary data are collected, making it suitable for a wide variety of power analyses in ST studies. We also provide a user-friendly, program-free web application (https://lanshui.shinyapps.io/PoweREST/), allowing users to interactively calculate and visualize the study power along with relevant the parameters.

## 1 Introduction

The recent progress in Spatial Transcriptomics (ST) enabled high-throughput measurements of transcriptomics while preserving spatial information about the tissue context (1). It has facilitated biological research in numerous domains, such as developmental biology, oncology, and neuroscience (2, 3). With the incorporation of both transcriptomics and spatial data (4, 5), ST data provide exciting opportunities to investigate human tissues from different angles, such as identifying the detailed tissue architecture, exploring domain-specific cell-cell interaction, and detecting differentially expressed genes (DEGs) between different regions (4, 6). Among these topics, DEG detection serves as one of the most fundamental problem. After the tissue structures are identified using pathological or computational approaches, DEG detection helps explain the heterogeneity across different tissue regions. The DEGs can serve as potential druggable targets for treatment and diagnosis (7–10).

Although ST technology has significantly advanced transcriptomic studies, the high cost of current ST profiling platforms limits the application of ST technology in large-scale studies (11). Several key experimental factors can affect signal generation in ST data, including the choice of tissue area, the number and size of the region of interest (ROI), and the number of spots. Recent studies have provided insights upon how the number and size of ROI can impact the statistical power but the application is restricted to cell type detection and cell-cell communication (12, 13). Due to the importance of DEG detection, the associated power analysis was well developed for bulk RNA-seq and single cell RNA-seq (scRNA-seq) experiments (6). However, there is a limited amount of research in the literature regarding the power calculation for detecting DEGs using ST samples. Consequently, there is often a pressing need to optimally utilize the available resources to reach a good statistical power.

In transcriptomic studies, the power is usually influenced by parameters like the desired error rate, the magnitude of the experimental effect of interest (effect size), and the sample size, which can be either the number of biological replicates or the number of cells/spots. In the case of detecting DEGs using bulk RNA-seq, the effect size is a gene’s mean expression ratio (fold change) across two experimental conditions. Furthermore, since bulk RNA-seq DEGs analysis usually involves multiple genes, multiple comparison issue needs to be addressed with Bonferroni adjustment or false discovery rate (FDR) (14). For scRNA-seq studies which have cell-level information available, DEG analysis can further focus on DEGs between different conditions for a specific cell type or DEGs that show differential expression across various cell types while under the same experimental condition. Therefore, more factors can influence the study power. Apart from effect size, number of biological replicas and multiple testing methods, number of cells and cell type proportion should also be taken into considerations (15, 16).

The additional coordinates information in ST data makes the power estimation for DEGs analysis more complicated than that of bulk RNA-seq and scRNA-seq. To the best of our knowledge, only one recent publication dedicated on the power analysis for a NanoString GeoMx ST experiment to study non-alcoholic fatty liver disease (17). Their method is not applicable to other popular ST platforms such as 10X Visium (18, 19). That is because GeoMx supports freeform ROIs which can be drawn to collect probes from any given region within the dimensions of 5-650 μm and 10X Visium measures gene expression in pre-determined sequenceable spot sizes of 55 μm with 100 μm spacing between spot centres. Therefore, the power of GeoMx experiments is influenced by the ROI’s shape and size while experiments based on 10X Visium is guided by the number of spots and specimens. Furthermore, (17) provides no accessible tool for non-coders which limits its wide application. Actually, so far there has been no accessible technique such as a software tool for power estimation upon ST experiments.

To address the aforementioned challenges, we present a novel **PoweR E**stimation tool for **ST** data, PoweREST (https://github.com/lanshui98/PoweREST) with a Shiny app (https://lanshui.shinyapps.io/PoweREST/). This tool enables choosing the optimal 10X Visium ST experimental design for DEGs detection between two conditions. Unlike most power calculation methods that simply assume the gene expression follows Poisson or negative binomial distributions (20, 21), PoweREST proposes a non-parametric statistical power evaluation framework based on in silico bootstrap datasets that resemble the real ST data. Moreover, our method employs the penalized spline (P-spline) technique (22) and XGBoost (23) under constraints to ensure the monotonic relationship between power and other parameters. Such monotone-respecting property was not considered by the power estimation method upon NanoString GeoMx ST data (17). These algorithmic and statistical advantages with robust results across different ROIs and different tissue samples support PoweREST as a practical tool for accurate power estimation of DEGs detection upon ST data.

## 2 Methods

### 2.1 PoweREST analytical framework

PoweREST evaluates how experimental design affects power and help identify an optimal sample size selection. PoweREST uses a nonparametric framework and simulate different experimental scenarios based on the real Visium ST dataset to fully accommodate the complexity of ST data. The methods consist of four steps: (i) Bootstrap resampling the spots within the ROI; (ii) Differentially expression (DE) analysis upon the resampled subjects; (iii) Estimation of the power based on the adjusted p-values for multiple testing; (iv) Monotonic estimation of the statistical power surface using P-splines with XGBoost as a remedy. The schematic overview of a PoweREST workflow with and without the ST dataset available is shown in Figure 1A.

**Fig. 1.**
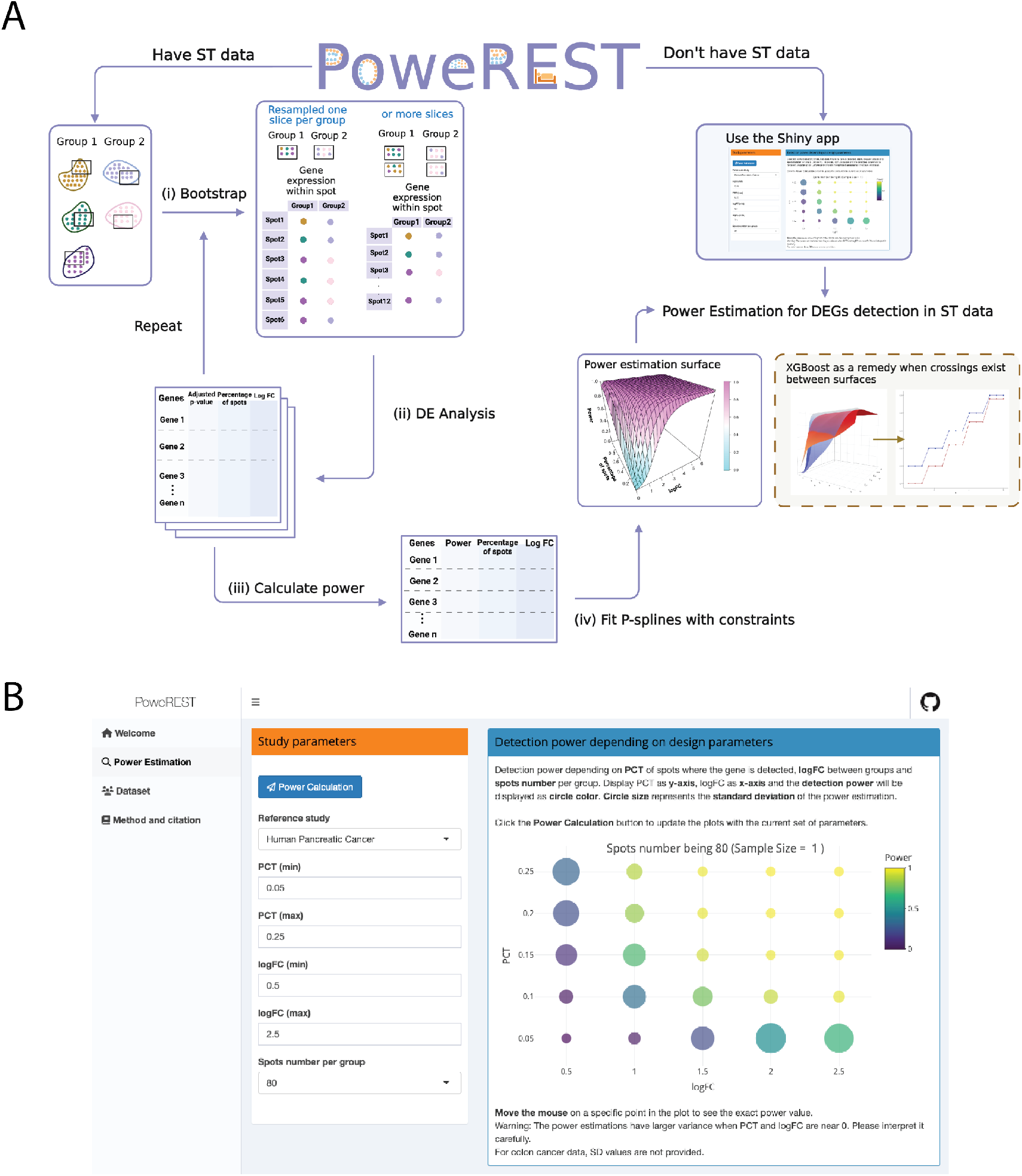
Schema of the proposed method. (A) Schematic overview of PoweREST workflow. When a preliminary cohort of ST data is available, PoweREST performs the power calculation based on bootstrapping (left and bottom) and P-splines (bottom right). When preliminary data is not available, an R shiny application was provided with power estimation results based on the datasets from two cancer studies (up right). (Created with BioRender.com) (B) A screen shot of the R shiny application. The shiny app provides straightforward power estimation for DEGs of given log fold changes and expression rates.

Users can apply PoweREST in two ways. When preliminary Visium ST data are available, users are welcomed to use the preliminary data as input and apply steps in Section 2.1.1 to Section 2.1.5 with our R software package to fit problem-specific power surface. Otherwise, users can directly provide the targeted effect sizes as parameters to the PoweREST shiny app (discussed in Section 2.2) and obtain power calculation based on our pre-trained models from two publicly available datasets described in Section 3.

#### 2.1.1 Data resampling

Let *C*_1_ and *C*_2_ be the true “population” ST datasets under condition 1 and 2, *c*_1_ be the random ST samples from condition 1, *c*_2_ be the random ST samples from condition 2. Assume there is an average of *n* spots selected in each slice across two conditions which are usually guided by H&E staining. Denote the average detection rate of a gene across two conditions as *π*_*g*_ and the log-fold change in average expression between two conditions as *β*_*g*_. The power that we wish to estimate is *P* = *P ower*(*n, π*_*g*_, *β*_*g*_, *α, N*) with the desired adjusted p-value being *α* and the target replicates (slices) number being *N* in each group. Since *n* is usually a fixed value in one experiment, for the rest of the paper, we view slices number *N* as the sample size and treat it as the influencing factor to power rather than *n*.

PoweREST creates ST specimen replicates within that ROI by bootstrap resampling (24, 25). Specifically, PoweREST randomly draws spot-level gene expression with replacement from the sample data *c*_1_ and *c*_2_ to mimic the sampling process from the true “population” *C*_1_ and *C*_2_. Our method also has the flexibility to create different numbers of in-silico replicates for each experimental group to serve the power calculations. In this paper, for simplicity, we assume the number of replicates is equal between conditions.

#### 2.1.2 DE analysis

After generating the synthetic specimens through bootstrap resampling, our method implements the *FindMarkers* function from *Seurat* package for DE analysis (26). This function performs the DE analysis between two groups using the Wilcoxon Rank Sum test in the default setting. Derived by Noether (27), the approximate power formula of Wilcoxon test on X and Y with equal sample sizes *n*_*xy*_ can be expressed as:

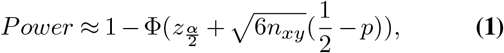

where *p* = *Prob*(*Y > X*), Φ is the CDF for the Standard Normal N(0,1) and 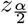 is an alpha level’s z-score for a two tailed test. When the Wilcoxon test is performed upon the synthetic specimens, *X* and *Y* would then be the gene expression in the first and second group. *n*_*xy*_ is the product of the spots number *n*, target replicates number *N*, and the percentage of spots detecting the gene *π*_*g*_. Intuitively, *Prob*(*Y > X*) can be influenced by the log fold-change *β*_*g*_. Thus, from the formula, we can infer that the study power grows with a higher absolute value of *β*_*g*_ or a larger expression rate. Such tendency can also be inferred from the real data (Supplementary Figures S1 and S5). However, the actual power formula is hard to be established without further assumptions placed upon the gene expression data. This is the reason why we choose non-parametric methods to estimate the power.

By applying the *FindMarkers* function to the resampled in-silico replicates, we can estimate the log fold-change *β*_*g*_ of the average expression between the two groups, the percentages of spots where the gene is detected in the first and second group. Additionally, we obtain the average value as the estimation for *π*_*g*_ and the adjusted p-value *α*_*g*_, which by default is based on Bonferroni correction. According to *Seurat* (26), other correction methods are not recommended, since *FindMarkers* pre-filters genes which already reduces the number of tests performed. These results are recorded and used for power assessment in Section 2.1.3.

#### 2.1.3 Power generation

In order to estimate the statistical power, the previous two steps (i) bootstrap sampling and (ii) DE analysis are repeated for many times, say 100 times. Within every repetitions *i*, the genes are considered to be DEGs when the adjusted p-value *α*_*gi*_ is less than *α*. The power of DEG detection is obtained by:

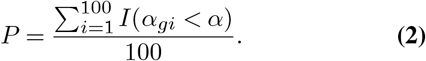

#### 2.1.4 P-splines fitting

After the previous three steps, the *P* values are derived under different combinations of (*N, π*_*g*_, *β*_*g*_) values. And the number of power values we can get is determined by the number of genes measured from the samples which is usually over thousands. In order to estimate the power under a new combination of (*π*_*g*_, *β*_*g*_) values with sample size *N*, PoweREST utilizes two-dimensional P-splines with monotonic constrains to fit a power surface.

When the sample size *N* is fixed, it is natural to assume that *P* increases as any one of (*π*_*g*_, |*β*_*g*_|) increases where |*β*_*g*_| is the absolute value of *β*_*g*_. In this way, unconstrained nonparametric models might be too flexible and give implausible or un-interpretable results. Our method utilized shape constrained additive models (*SCAM*) (28, 29) to fit the power surface while preserving the monotonicity between *P* and both (*π*_*g*_, |*β*_*g*_|) parameters:

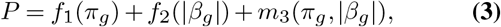

where *f*_1_ and *f*_2_ represent smooth “main effects” and *m*_3_ is a smooth “interaction”. Specifically, for a univariate smooth spline function f:

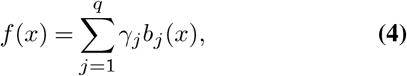

where *q* is the number of basis functions, *b*_*j*_ are B-splines basis functions, and *γ*_*j*_’s are unknown coefficients. A smoothing penalty and a shape constraint are imposed upon *γ*_*j*_ to control the “wiggliness” of *f* while ensuring the monotonic relationship between *f* and *x. SCAM* uses quadratic splines and applies Newton-Raphson method to maximize the penalized likelihood for *γ*_*j*_’s estimations. The estimations are robust to the choice of *q* when shape constraints are employed (30). The statistical expression and derivation of the bivariate P-splines *m* under double penalties and double monotonicity can be found in (29).

#### 2.1.5 XGBoost as a remedy

P-splines with 2-dimensional monotonic constraints ensure the monotone relationships between power and any one of (*π*_*g*_, |*β*_*g*_|), but the monotonic relationship between power and sample size *N* is not respected. Currently, a robust software for P-splines under 3-dimensional monotonic constraints is not readily available. In practice, we found the estimated power values adhere the monotonicity with sample size when (*π*_*g*_, *β*_*g*_) are large but this relationship breaks down in some cases when both parameters are close to 0. To address this, we propose a remedy which employs XGBoost (23) to impose 3-dimensional monotonic constraints upon all the three predictors: *π*_*g*_, |*β*_*g*_| and *N* for estimating power values when (*π*_*g*_, *β*_*g*_) are small.

XGBoost approaches the prediction problem by decision tree ensembles which sum up the decision values of multiple trees together for a final decision. The tree structural is trained through an additive strategy: fix what have been learned, and select the new leaf which optimizes the objective at a time. The monotonic constraints are achieved by that at every step, the algorithm will abandon a candidate split if it causes a non-monotonic relationship. However, as it essentially treats the prediction problem as a kind of decision making or classification, it fits visually a step function rather than a smoothing curve. Therefore, using XGBoost to fit the entire power surface can be crude. We recommend that users begin with P-splines and carefully examine the resulting fits. When power surfaces for different sample sizes intersect, consider applying XGBoost specifically within the regions where these crossings occur. But don’t mix the results from P-splines and XGBoost together, since they are fitted under different algorithms. The details about how the monotonic constraints are preserved at the algorithm level can be found in (23) and its GitHub repository.

### 2.2 Implementation of software package and shiny app

We implemented the proposed methods in an open-source R package PoweREST. A tutorial is distributed on the GitHub pages, which contains detailed instruction and examples of using the package and interpreting the results. To facilitate the application by users who are not familiar with R coding, we also create an online, interactive, program-free web application using R Shiny. As shown in Figure 1B, users can select the preferred tissue type with targeted parameter values of the ST experiments and the study power can be generated by clicking the “calculate” button. The Shiny app currently only incorporates the results from P-splines, the results of XGBoost can be found on the GitHub pages.

## 3 Results

### 3.1 Power surface estimation with human intraductal papillary mucinous neoplasms (IPMN) data

The first dataset we examined was a publicly available 10X Visium dataset (GSE233254) from human IPMN tissues (10), which contains 13 specimens with 12,685 spots covering the specimens and up to 8,000 genes detected. The 13 specimens are divided into two categories: 6 specimens in the “high-risk” (HR) category and 7 specimens in the “low-grade” (LG) category. IPMNs are bona fide precursor lesions of pancreatic ductal adenocarcinoma. Clinically, HR lesions are those mandate surgical resection, with or without an associated invasive cancer (10). In order to reveal area-specific changes in DEGs between two groups, the researchers have annotated the spots overlapping with the neoplastic epithelium (“epilesional”, n = 755); the immediately adjacent microenvironment, corresponding to two layers of spots (∼ 200 *μ*m) surrounding the lining epithelium (“juxtalesional”, n = 1,142); and an additional two layers of spots located further distal to the juxtalesional region (“perilesional”, n = 1,030) based on the H&E staining. Figure 2A-B show two representative slices of such histologically direct spot annotation. Within each region, we resampled the spots to create simulated samples within that region and performed the proposed power estimation method.

**Fig. 2.**
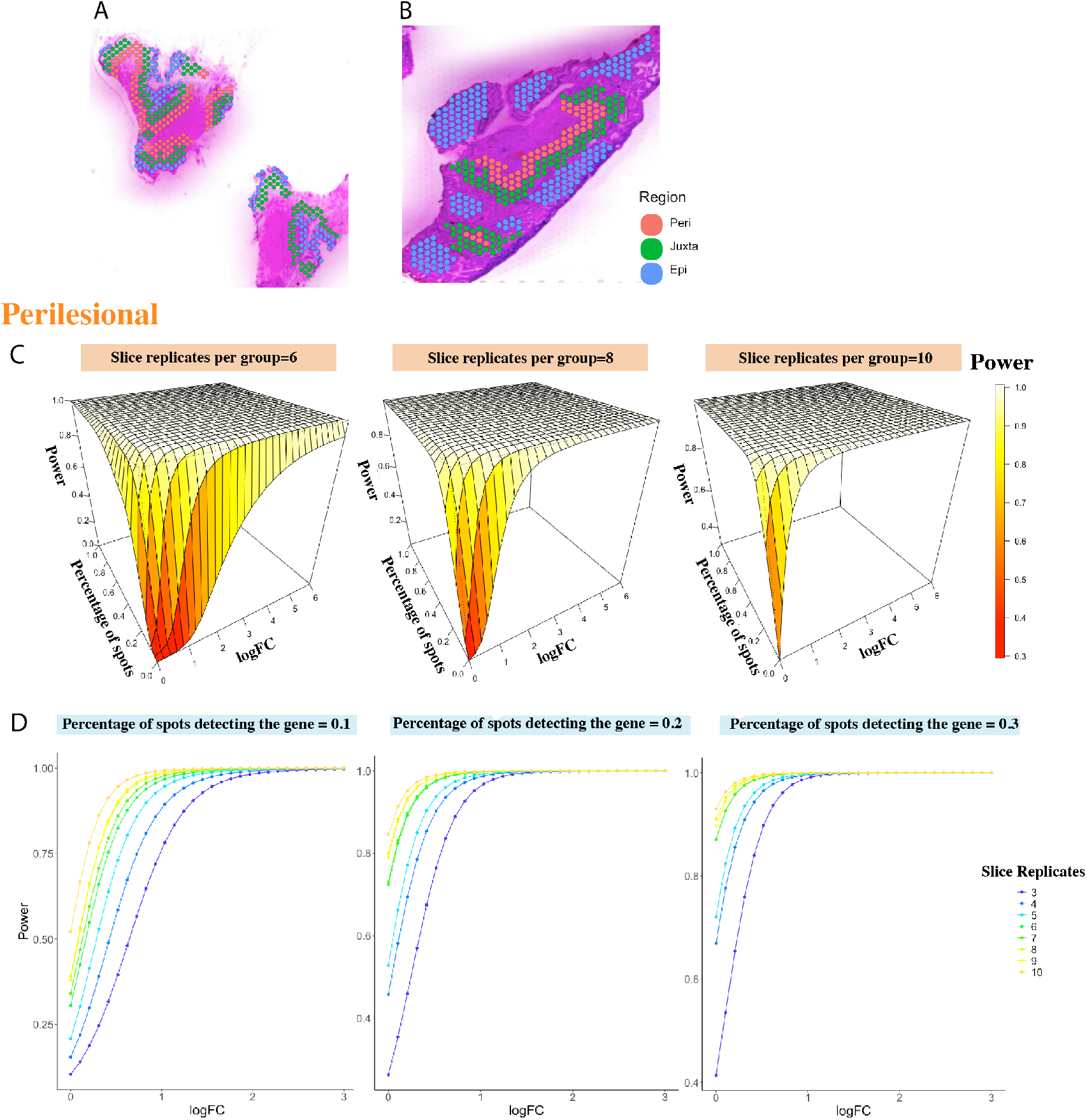
Power estimation based on the IPMN dataset. Histologically annotated epilesional, juxtalesional, and perilesional areas are shown in (A) an LG sample and (B) a HR sample. (C) The fitted power surfaces for sample size being 6, 8, 10 per group, of DE analysis within perilesional areas. The power was fitted under the constrains that it monotonely increases with the percentage of spots where the gene can be detected and the log fold change. (D) The relationships between the power and log fold change when the percentage of spots detecting the gene equals 0.1, 0.2, 0.3. While we didn’t force the monotone relationship between sample size and power, their monotonic relationship is still respected from the fitted results.

#### 3.1.1 Power of DE analysis in perilesional area

For a total of 1,030 spots annotated as perilesional area, we have 540 spots from HR samples and 490 spots from LR samples. We resampled 240, 320, .., 720, 800 spots 100 times from HR and the same number of spots from LG to mimic the regional specimens of 3, 4, … 10 replicates under each condition. We resample the spots at an increment of 80 for the reason that roughly there are 80 spots per specimen at each region. Through the PoweREST pipeline, the power surfaces were fitted smoothly for different combinations of logFC and percentage values while maintaining the monotonic relationships. The fitted surfaces for three selected replicate values were shown in Figure 2C. Although the power surfaces were estimated separately for each replicate value, the foreseeable monotonic relationship between power and number of replicates per group remained (Figure 2D). As a comparison, we used XGBoost to model power values, based on raw power values where the logFC was below 1 and show the results in Supplementary Figure S4.

#### 3.1.2 Validation in juxtalesional and epilesional areas

In order to validate the proposed method across different ROIs from the same tissue, we repeated the analysis in the other two areas. For the juxtalesional area, we have in total 1,142 spots where 568 spots from HR samples and 574 spots from LG. And among the 755 spots annotated as the epilesional area, 441 spots are HR and 314 spots are LG. To obtain comparable results across different areas, the spots were still resampled at an increment of 80. Again, PoweREST estimates the power under different values of parameters using P-splines. The power results for juxtalesional and epilesional areas are presented in Figure 3A-B and Supplementary Figure S2 which are similar to the power surfaces we obtained for the perilesional area. The relative difference in the power results between two different areas was calculated (Figure 3C, Supplementary Figure S3), from which we found that that the differences are minimal and only exist in regions where both the logFC and the percentage of expressed spots are low. The relative difference between juxtalesional and perilesional areas is from 1.8e-10 to 1.2, and from 9.5e-09 to 7.4 for the relative difference between epilesional and perilesional areas, where 7.4 is the relative difference when logFC and percentage of spots with gene expressed being 0s and the power estimation values are 0.3 and 0.04 respectively.

**Fig. 3.**
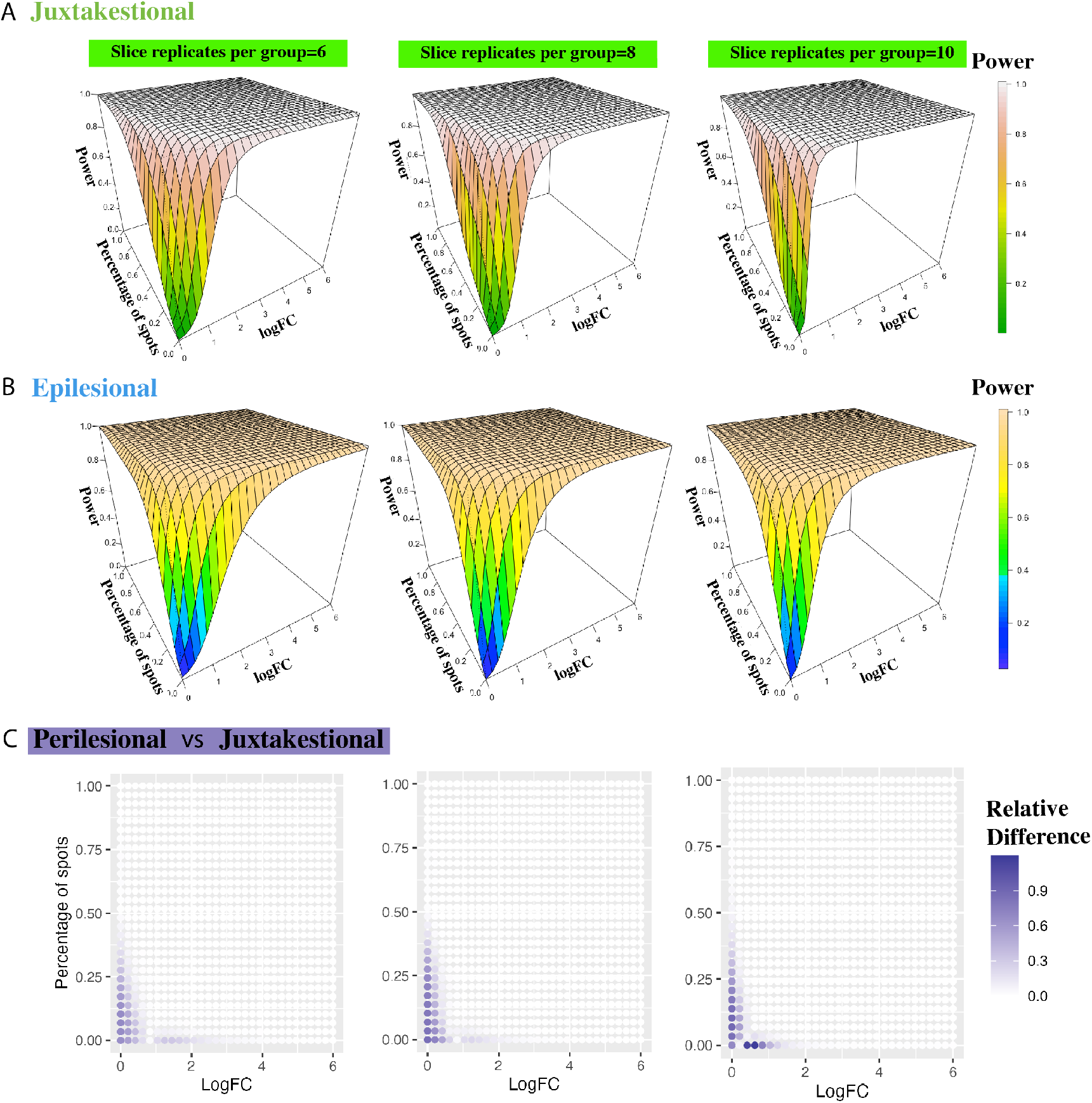
Validation upon the other two areas from IPMN dataset. The fitted power surfaces for slice replicate number being 6, 8, 10 per group, of DE analysis within juxtalesional areas (A) and epilesional areas (B). (C) The relative difference between the estimated power surfaces from perilesional areas and juxtalesional areas, when the number of replicates per group is 6, 8, 10.

These observations suggest that PoweREST robustly estimate statistical power for DEG detection across different functional regions in the IPMN samples.

### 3.2 Power surface estimation with human colorectal cancer (CRC) data

The second dataset we used to apply our methodology was a human CRC dataset measured by 10X Genomics (31). The dataset contains 31 human colonic specimens with around 17,000 gene detected, from which we selected 7 tissue samples diagnosed as microsatellite unstable (MSI-H) and 6 tissue samples with microsatellite stable (MSS). As is described in the paper (31) MSI-H CRCs are more immunogenic than their conventional adenoma MSS CRCs on average. Therefore, it is meaningful to figure out DEGs between the two disease conditions. We have included the used data in the GitHub site and shiny app.

#### 3.2.1 Power surface estimated by P-splines

As shown in Figure 4B-C, two areas (carcinoma and carcinoma border) were annotated by pathologists based on their H&E staining information. We focused our analysis upon the carcinoma border which approximately contains 500 spots per slice. We resampled 500, 1000, .., 4500, 5000 spots 100 times from MSS and the same number of spots from MSI-H to mimic the regional specimens of 1, 2, … 10. The power estimation results are shown in Figure 4A. Since the number of spots within one slice is larger than that of IPMN samples, a higher power value can be achieved with the same number of slice replicates and other two variables, logFC and percentage of spots. Figure 4D shows there exist crossings between power surfaces of different sample size value but occur only when logFC is small and the estimation may not be numerically stable.

**Fig. 4.**
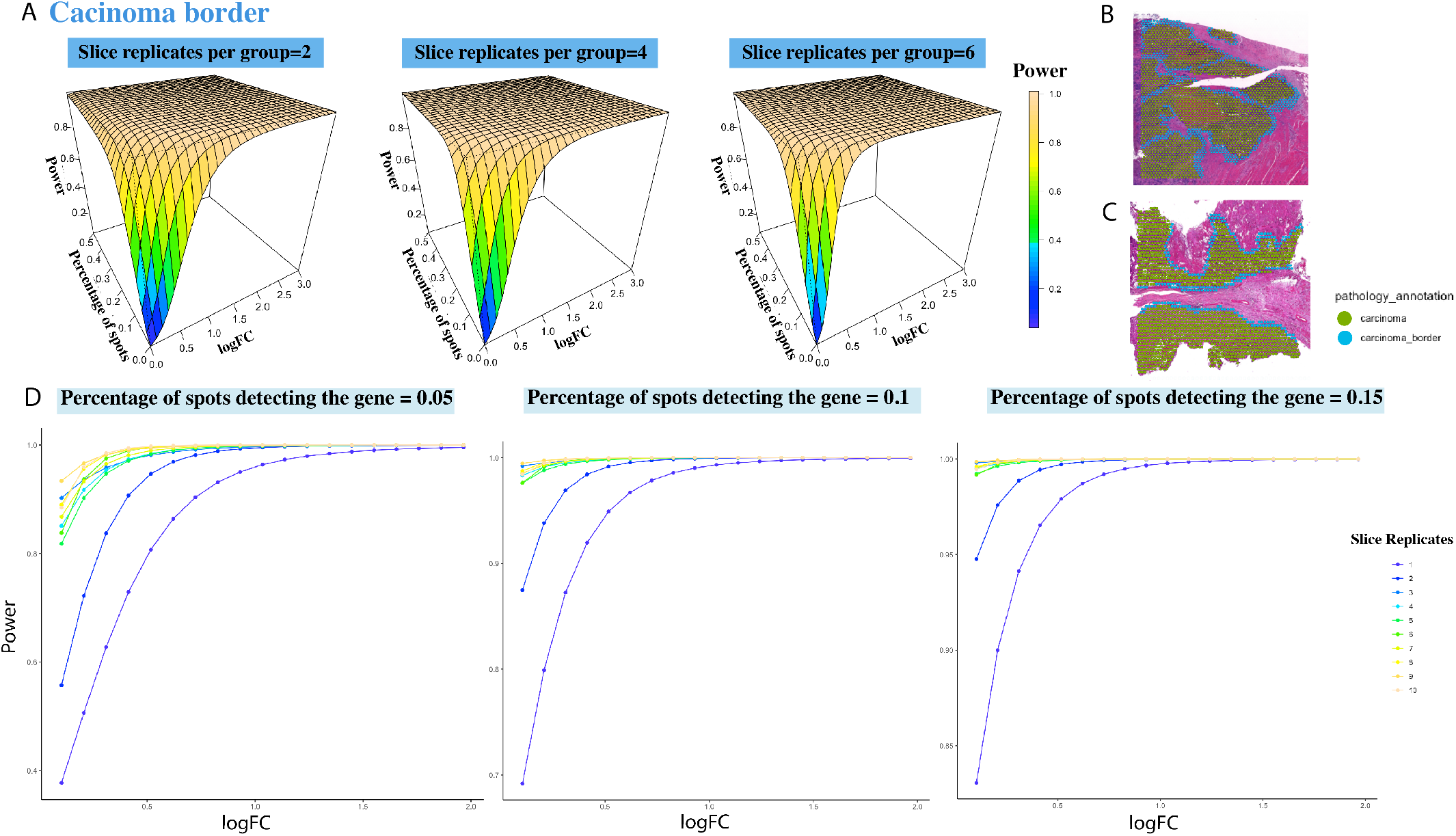
Power estimation based on the CRC dataset. (A) The fitted power surfaces using P-splines for DE analysis within the carcinoma border, under the slice replicates per group being 2, 4, 6. Histologically annotated carcinoma and carcinoma border areas in a MSS sample (B) and a MSI-H sample (C). (D) The relationship between the power and logFC under slice replicates from 1 to 10, for the percentage of spots detecting the gene being 0.05, 0.1, 0.15.

#### 3.2.2 Local power estimation by XGBoost

As a solution, XGBoost with 3-dimensional monotonic constraints can fit the power values when parameters are small, which deals with the crossings between power surfaces fitted by P-splines. The estimated power values for logFC less than 1.5 and expression rates below 0.15, as derived using XGBoost, are shown in Figure 5. These estimates resemble step functions with no intersections among the surfaces of difference sample size. However, when we tried to implemented XGBoost across a broader range of values for logFC and expression rates, the power estimation proved ineffective (Supplementary Figure S6). This may be because in this example, the power values increased rapidly with even a minor increase in parameter values, which hindered accurate power assessment by XGBoost’s classification strategy. By contrast, quadratic spines implemented in *SCAM* are capable of catching such patterns. Therefore, XGBoost is recommended for local power value estimations when crossings occur in regions with parameters being of specific interest.

**Fig. 5.**
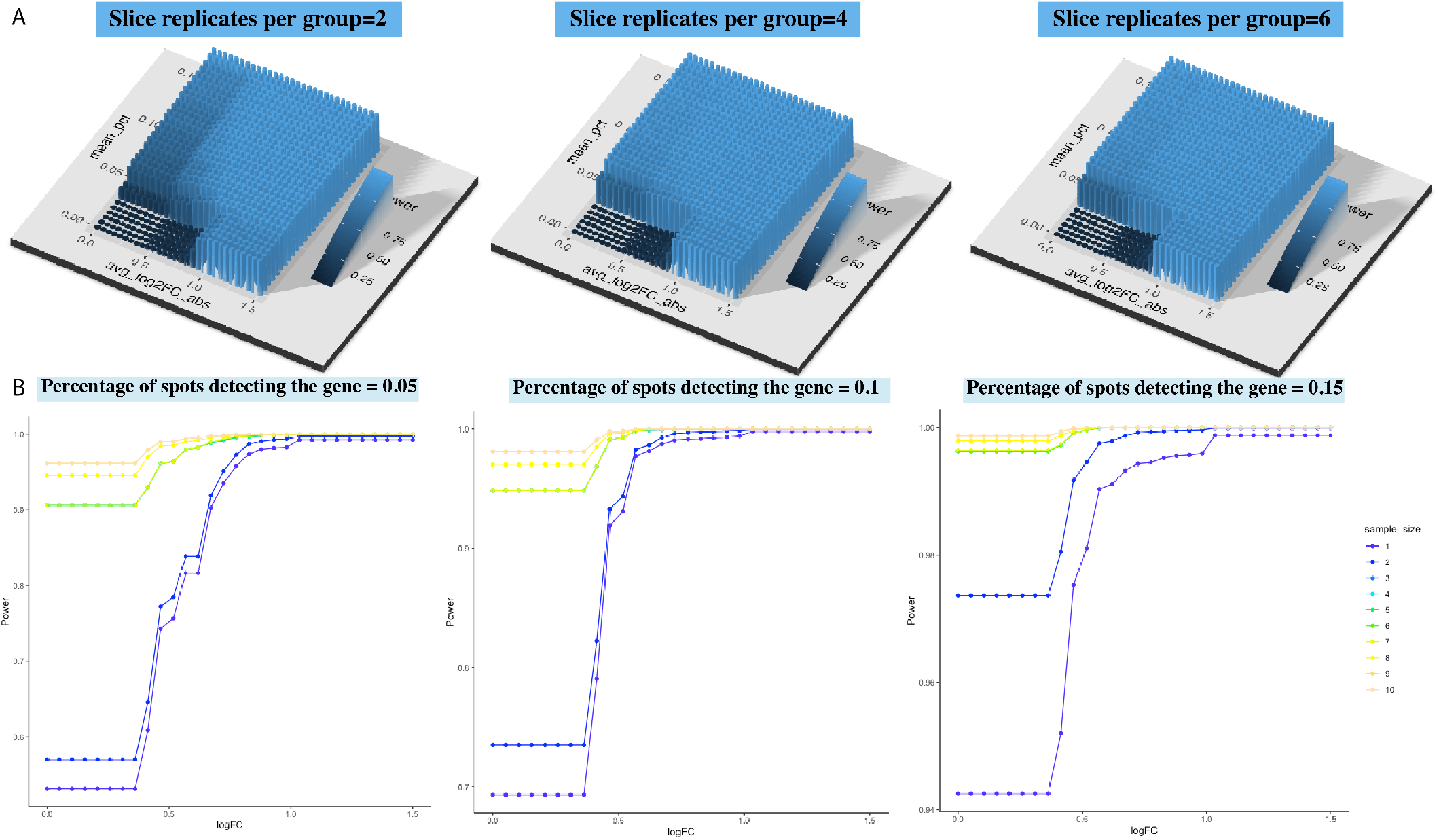
Power estimation based on the CRC dataset. The power is fitted by XGBoost under the constrains that it monotonely increases with the percentage of expressed spots, the log fold change and the number of slice replicates and the estimation is based on power values where logFC ≤1.5 and percentage of expressed spots ≤0.15. (A) The fitted power surfaces for DE analysis within the carcinoma border, under the slice replicates per group being 2, 4, 6. (B) The relationship between the power and logFC under slice replicates from 1 to 10, for the percentage of spots detecting the gene being 0.05, 0.1, 0.15.

## 4 Discussion

We have developed PoweREST, a novel and flexible method of power estimation, for detecting DEGs between two groups using ST data. One of the key considerations when designing a biomedical experiment is determining the appropriate sample size to ensure adequate statistical power. While various methods have been developed for bulk RNA-seq and single-cell RNA-seq, power estimation for ST data remains underexplored due to the challenges posed by its complex data structure and the integration of spatial information. (6, 14, 15). The present study introduces a fully non-parametric pipeline to depict the power for experiments with 10X Visium ST data. Unlike Baker et al. (13) who devised methods for tunable “in silico” tissue (IST) generation, our method used bootstrap to generate real data-like ST samples in a more transparent way. (24) The estimated power values are then calculated by repeating the DE analysis upon the generated ST samples. That our method is fully non-parametric is also illustrated by utilizing the shape-constrained P-splines (30) to fit the power surface along values of the logFC and the gene detection rate among spots. As an intrinsic requirement, two-dimensional monotone constrains were imposed on the P-splines which respects the monotonicity of the estimated power surface with the increase of logFC and expression percentage. XGBoost was proposed as a remedy to crossings between power surfaces.

As presented in Section 2.1.2, Noether (27) derived a formula to approximate the study power in the Wilcoxon Rank Sum test using *Prob*(*Y* > *X*). However, there is no deterministic relationship between *Prob*(*Y* > *X*) and effect size unless additional distribution assumptions are put upon the ST gene set. Additionally, the sparsity in the ST data leads to large amount of tied observations which violates the assumption of Noether’s formula (32). The formula based power calculation also did not consider the multiple testing issue (33). Compared with formula-based power calculation, the non-parametric modeling depends on least amount of assumptions and thus is more generalizable to the complex power estimation with ST data. One disadvantage of the bootstrap-based approach is the long computation time. As solutions, we provide the pilot results in the R package and R shiny app upon different tissue types. Similiar power results were shown across different areas within the same tissue type. In the R package, we provide options to pre-filter the genes based on their minimum detection rate and at least X-fold difference to speed the process. We also include functions for power calculation focusing on those genes specified by users.

PoweREST utilizes the P-splines under two-dimensional monotone constraints. Such constraints ensure the monotone increasing relationships between power and logFC, and between power and detection rate of genes. However, the method doesn’t respect the monotone relationship between power and sample size. Instead, the method fits the power surfaces separately for each sample size value. In practice, it was observed that the monotonic relationship between estimated power and sample size is respected by the method in some cases. However, in other situations, this monotonic relationship is violated when the parameters approach 0. Currently, a robust software for P-splines under 3-dimensional monotonic constraints is not readily available. To address this, we proposed a solution using XGBoost, a machine learning technique that is capable of imposing three or more monotonicity constraints on the predictors. It essentially treats the prediction as a classification problem which fits visually a step function rather than a smooth curve (23). However, the method can be too crude to estimate the entire power surface. Therefore, we recommend that users initially employ P-splines and evaluate the resulting fits. When crossings occur at the targeted parameters, apply XGBoost, but restrict its use to the regions where these crossings arise.

Though non-parametric method makes minimal assumptions about the underlying distribution of the data being studied, it may sometimes lead to relatively larger residuals compared to parametric models. Such observations can be found in papers like (34), where they compared the residuals from P-splines and those from a Poisson distribution. Larger residuals with P-splines may be results of the emphasis on smoothness at the expense of fit or the presence of noise in the data that the model smooths over. Therefore, check residuals plots to diagnose where the model might be underperforming and interpret those results carefully. The function to create diagnosis plots are included in the R package.

## Author contributions statement

Z.L. and L.L. conceived the idea. L.S. performed the analyses and constructed the software package. Y.Y. provided useful suggestions on model formulation. L.S., Z.L. and L.L. wrote the manuscript. All authors reviewed and approved the final manuscript.

## Acknowledgements

This work was funded in part by the Coordination and Data Management Center (CDMC) of the Translational and Basic Science Research in Early Lesions (TBEL) Program, which is supported by the National Cancer Institute grant U24CA274212. Diagrams were created with BioRender.com.

## Data availability

This study made use of publicly available datasets. These include IPMN dataset (https://www.ncbi.nlm.nih.gov/geo/query/acc.cgi?acc=GSE233254) and CRC dataset (https://osf.io/hftq2/).

