## Supplementary Files for "PoweREST: Statistical Power Estimation for Spatial Transcriptomics Experiments to Detect Differentially Expressed Genes Between Two Conditions"

### 1 List of Figures for IPMN data

#### 1.1 Figure S1: Power results for IPMN's perilesional area before smooth fitting.

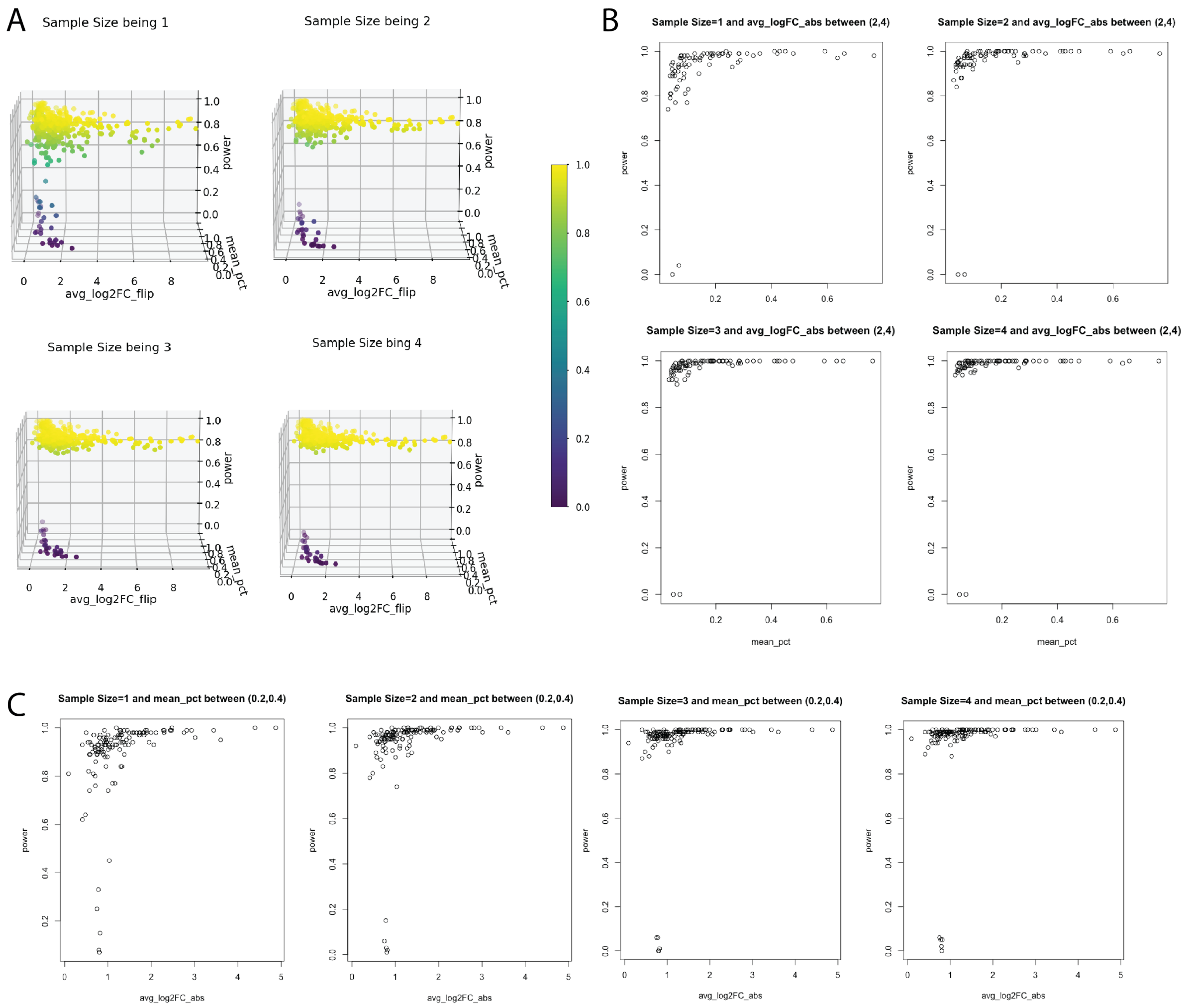

(A) The power values under biological replicates from 1 to 4. (B) Power values versus the percentage of expressed spots under biological replicates from 1 to 4, for the logFC between $(2, 4)$. (C) Power values versus logFC under biological replicates from 1 to 4, for the percentage of expressed spots between $(0.2, 0.4)$.

#### 1.2 Figure S2: The relationships between the power and log fold change for IPMN's juxtalesional and epilesional areas.

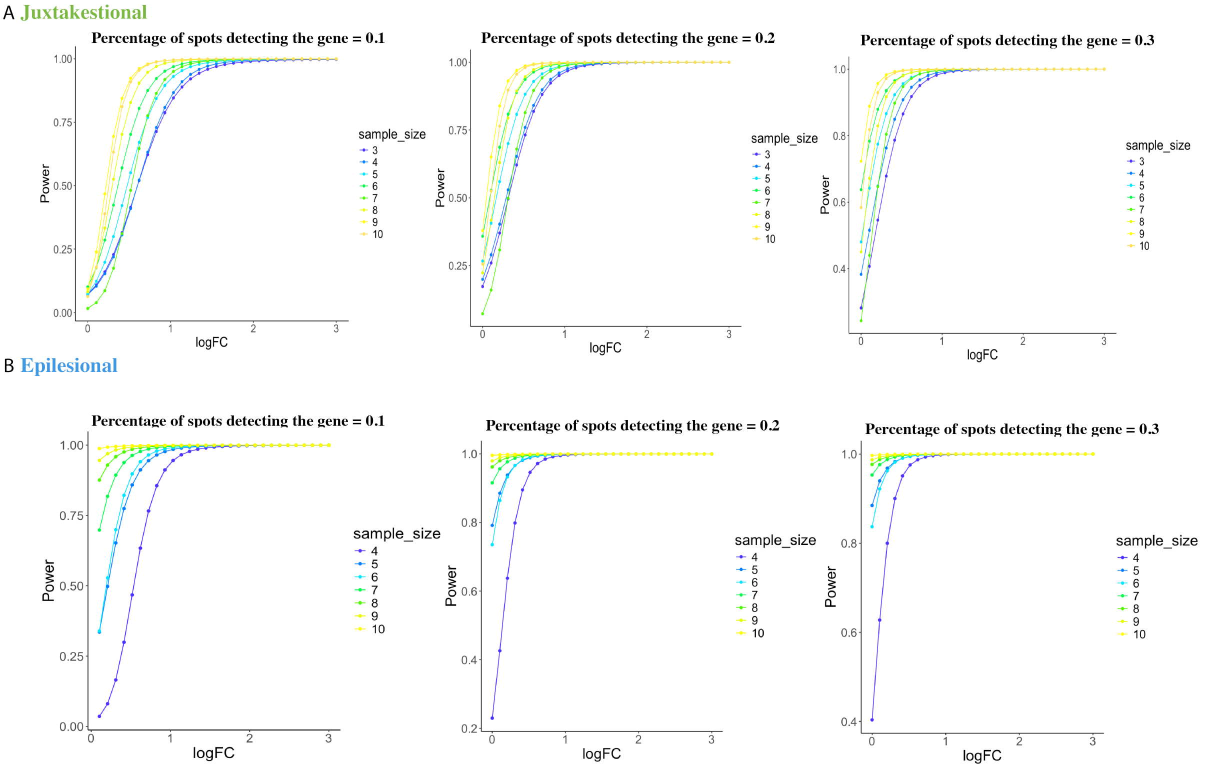

The relationships between the estimated power and log fold change when the percentage of spots detecting the gene equals $0.1, 0.2, 0.3$ for the juxtalesional areas (A) and epilesional areas (B).

#### 1.3 Figure S3: The relative difference between the estimated power surfaces from perilesional and epilesional areas.

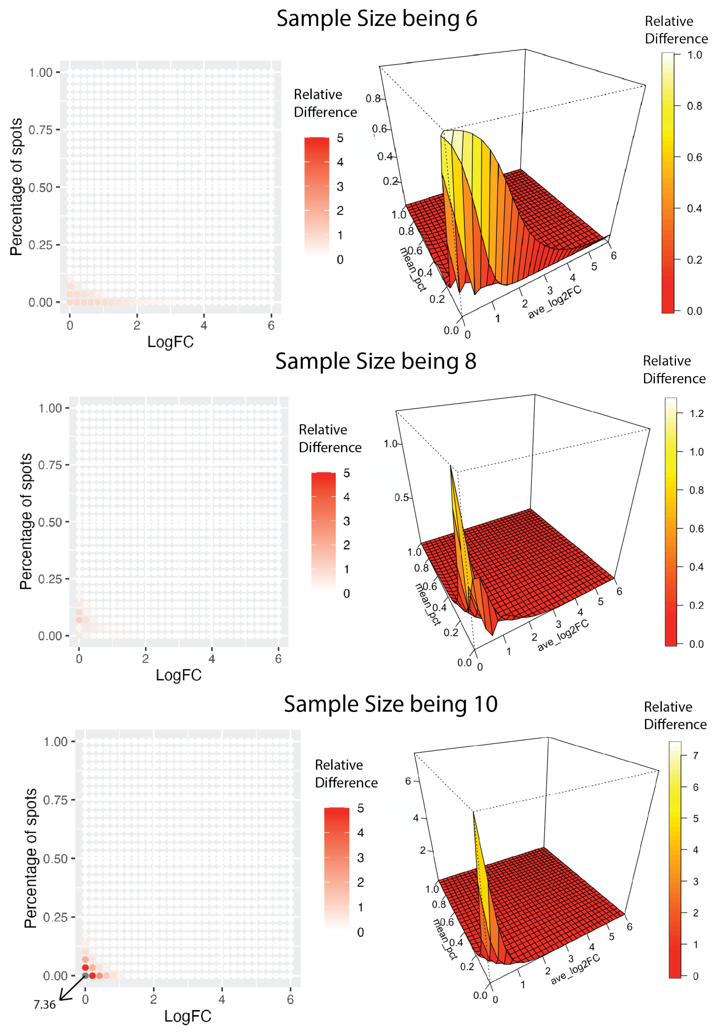

The relative difference between the estimated power surfaces from perilesional areas and juxtalesional areas, when the number of replicates per group is $6, 8, 10$.

#### 1.4 Figure S4: Power estimation of perilesional area based on XGBoost.

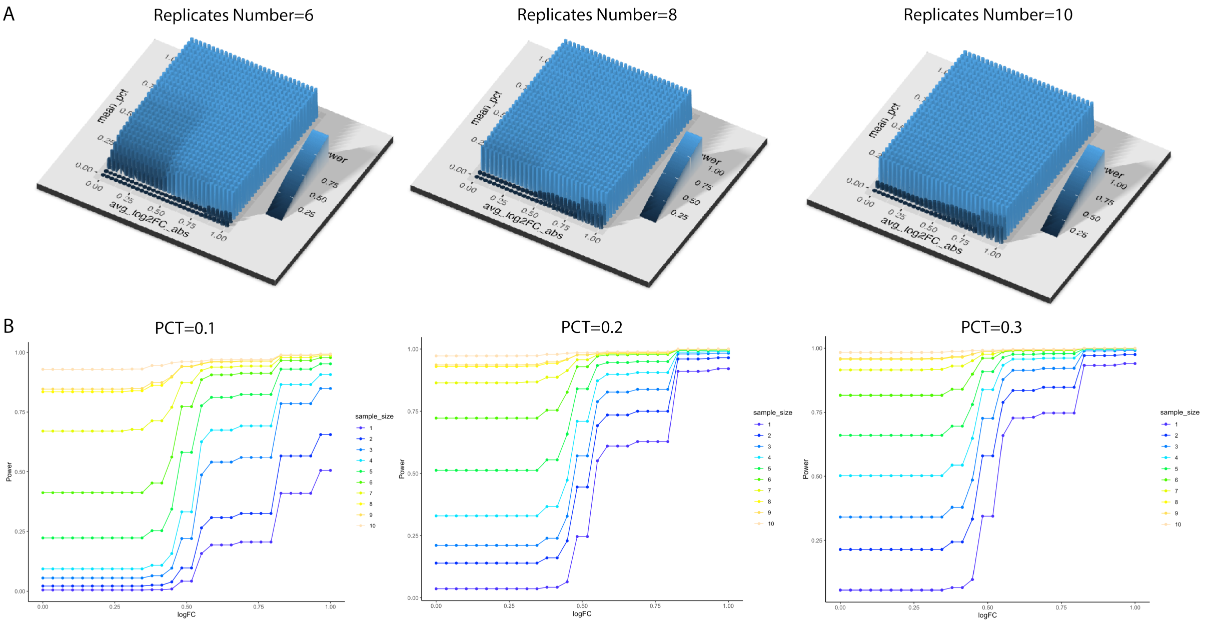

The estimation is based on raw power values where logFC is below 1. (A) The fitted power surface for sample size being 6, 8, 10 per group, of DE analysis within perilesional areas. The power is fitted by XGBoost under the constrains that it monotonely increases with the percentage expressed spots, the log fold change and the number of slice replicates. The results are comparable with Figure 2C in the main manuscript. (B) The fitted power versus logFC when the percentage of spots detecting the gene equals 0.1, 0.2, 0.3.

### 2 List of Figures for CRC data

#### 2.1 Figure S5: Power results for CRC data before smoothing fitting.

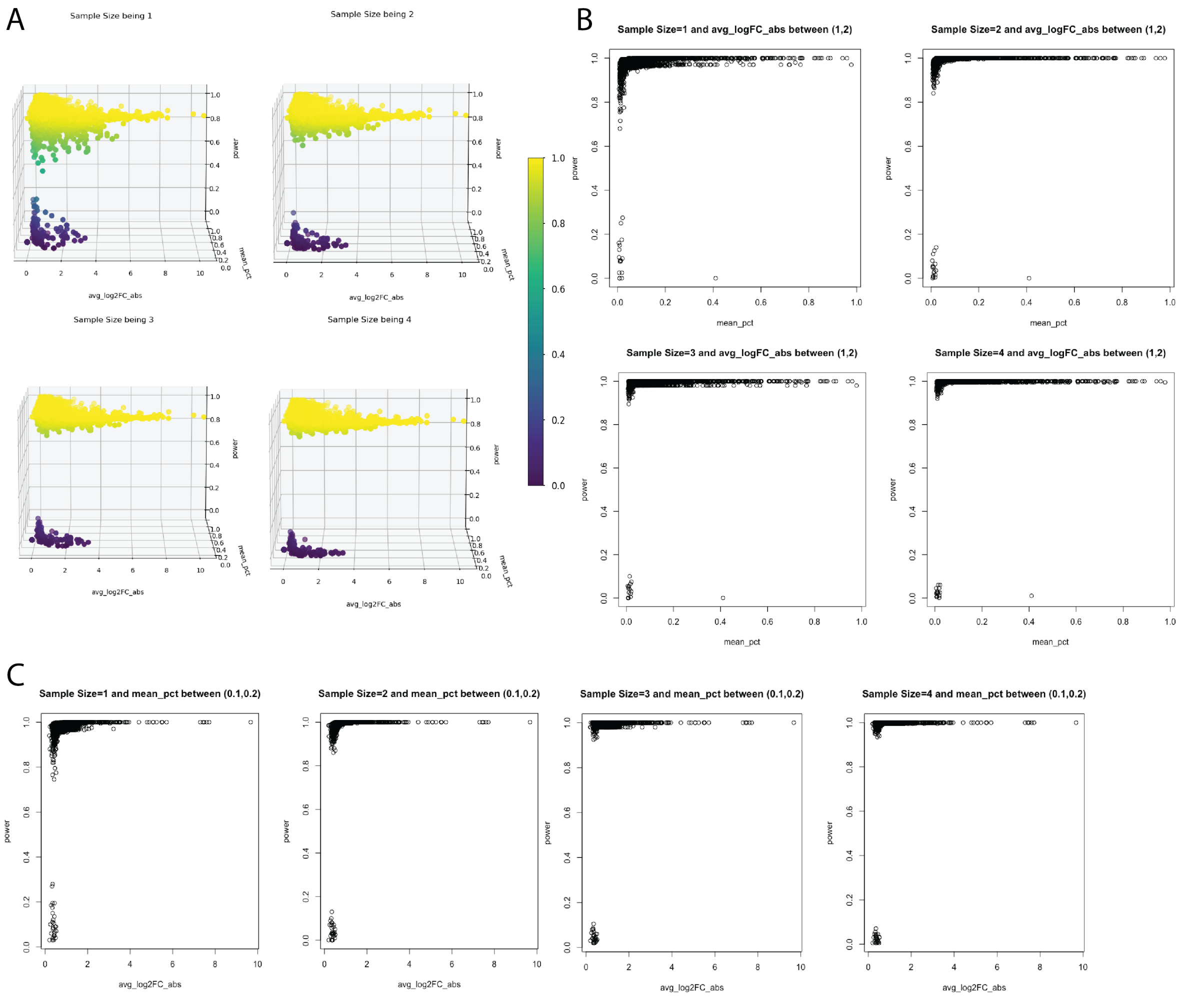

(A) The power values under biological replicates from 1 to 4. (B) Power values versus the percentage of expressed spots under biological replicates from 1 to 4, for the logFC between $(2, 4)$. (C) Power values versus logFC under biological replicates from 1 to 4, for the percentage of expressed spots between $(0.2, 0.4)$.

#### 2.2 Figure S6: Estimation upon the entire power surface using XGBoost.

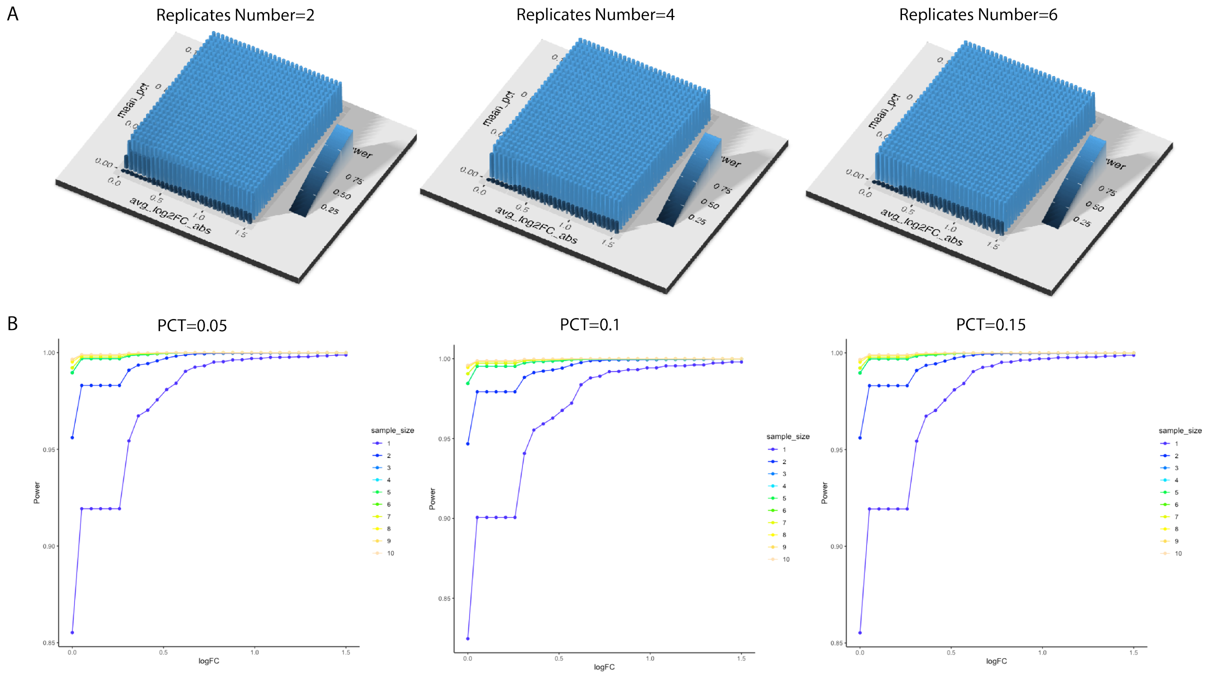

(A) The fitted power surfaces for DE analysis within the carcinoma border, under the slice replicates per group being $2, 4, 6$. (B) The relationship between the power and logFC under slice replicates from 1 to 10, for the percentage of spots detecting the gene being $0.05, 0.1, 0.15$. Compare with Figure 5 in the main manuscript where XGBoost was used to fit power values where logFC $\leq$1.5 and percentage of expressed spots $\leq$0.15, the estimations here are less precise due to the characteristics of XGBoost's algorithm.
